# Inhibition of TXNRD or SOD1 overcomes NRF2-mediated resistance to β-lapachone

**DOI:** 10.1101/849927

**Authors:** Laura Torrente, Nicolas Prieto, Aimee Falzone, David A. Boothman, Eric B. Haura, Gina M. DeNicola

## Abstract

Alterations in the NRF2/KEAP1 pathway result in the constitutive activation of NRF2, leading to the aberrant induction of antioxidant and detoxification enzymes, including NQO1. The NQO1 bioactivatable agent β-lapachone can target cells with high NQO1 expression but relies in the generation of reactive oxygen species (ROS), which are actively scavenged in cells with NRF2/KEAP1 mutations. However, whether NRF2/KEAP1 mutations influence the response to β-lapachone treatment remains unknown. To address this question, we assessed the cytotoxicity of β-lapachone in a panel of NSCLC cell lines bearing either wild-type or mutant KEAP1. We found that, despite overexpression of NQO1, KEAP1 mutant cells were resistant to β-lapachone due to enhanced detoxification of ROS, which prevented DNA damage and cell death. To evaluate whether specific inhibition of the NRF2-regulated antioxidant enzymes could abrogate resistance to β-lapachone, we systematically inhibited the four major antioxidant cellular systems using genetic and/or pharmacologic approaches. We demonstrated that inhibition of the thioredoxin-dependent system or copper-zinc superoxide dismutase (SOD1) could abrogate NRF2-mediated resistance to β-lapachone, while depletion of catalase or glutathione was ineffective. Interestingly, inhibition of SOD1 selectively sensitized KEAP1 mutant cells to β-lapachone exposure. Our results suggest that NRF2/KEAP1 mutational status might serve as a predictive biomarker for response to NQO1-bioactivatable quinones in patients. Further, our results suggest SOD1 inhibition may have potential utility in combination with other ROS inducers in patients with KEAP1/NRF2 mutations.

**Highlights:**

- Aberrant activation of NRF2 in non-small cell lung cancer promotes resistance to β-lapachone via the antioxidant defense.
- Inhibition of the thioredoxin-dependent system and superoxide dismutase 1 increase sensitivity to β-lapachone treatment.
- Mutations in the NRF2/KEAP1 pathway might serve as predictive biomarker for response to β-lapachone in patients.

## Introduction

It is estimated that 38% of lung squamous cell carcinomas (SCC) and 18% of lung adenocarcinomas (LuAD) harbor mutations in Nuclear factor erythroid 2-related factor 2 (NRF2), or its negative regulator Kelch-like ECH-associated protein 1 (KEAP1)^1–3^, making this pathway one of the most commonly mutated non-small cell lung cancer (NSCLC). The transcription factor NRF2 acts as the primary cellular barrier against the deleterious effects of oxidative stress by regulating the expression of cytoprotective genes. In healthy tissues, KEAP1 binds to and harnesses the activity of NRF2, thereby promoting NRF2 ubiquitination and destruction by the proteasome^4–6^. Loss-of-function mutations in KEAP1 and gain-of-function mutations in NRF2 found in NSCLC abolish this control and lead to constitutive NRF2 activity^1, 7–9^. Cancer cells that hijack NRF2 activity are equipped with a reinforced cytoprotective system through the induction of antioxidant and drug detoxification pathways, thereby rendering them resistant to oxidative stress and chemo/radio-therapy^10–12^.

High expression of the detoxification enzyme NAD(P)H:quinone oxidoreductase 1 (NQO1) is a distinct biomarker of NRF2/KEAP1 mutant NSCLC tumors. NQO1 is a cytosolic flavoprotein that catalyzes the two-electron reduction of quinones into hydroquinones in an effort to hamper oxidative cycling of these compounds^13, 14^. Although NQO1-dependent reduction of quinones has been historically defined as a major detoxification mechanism, a number of quinones induce toxicity following NQO1 reduction^15–19^. The mechanism behind this paradox relies on the chemical properties of the hydroquinone forms. Unstable hydroquinones can be reoxidized to the original quinone by molecular oxygen, which leads to the formation of superoxide radicals. As the parent quinone is regenerated, the cycle continues, which amplifies the generation of superoxide radicals, initiating a cascade of reactive oxygen species (ROS).

The ability of NQO1 to generate cytotoxic hydroquinones has been utilized as a strategy to target cancer cells with high NQO1 levels. To date, β-lapachone and its derivatives are the most studied NQO1-bioactivatable quinones, and the molecular mechanisms by which they promote cytotoxicity have been thoroughly characterized^20–24^ (Figure 1A). NQO1 has been proposed as a target for NSCLC therapy, as it is overexpressed in lung tumors but not in adjacent normal tissues^25, 26^. Thus, systemic delivery of β-lapachone would spare healthy lung tissue while inducing robust cytotoxicity in tumor cells. Remarkably, a large fraction of NSCLC with high NQO1 also harbor sustained NRF2 activation, which in turn could hinder the cytotoxic effects of β-lapachone through the active scavenging of ROS. Therefore, although high levels of NQO1 could be exploited with a therapeutic intent in NRF2/KEAP1 mutant cancer cells, it is unclear whether actions of NRF2 could limit β-lapachone efficacy. In this study, we aim to clarify whether NQO1 represents a druggable strategy for NRF2/KEAP1 mutant NSCLC or, conversely, these alterations promote resistance to β-lapachone.

**Figure 1.**
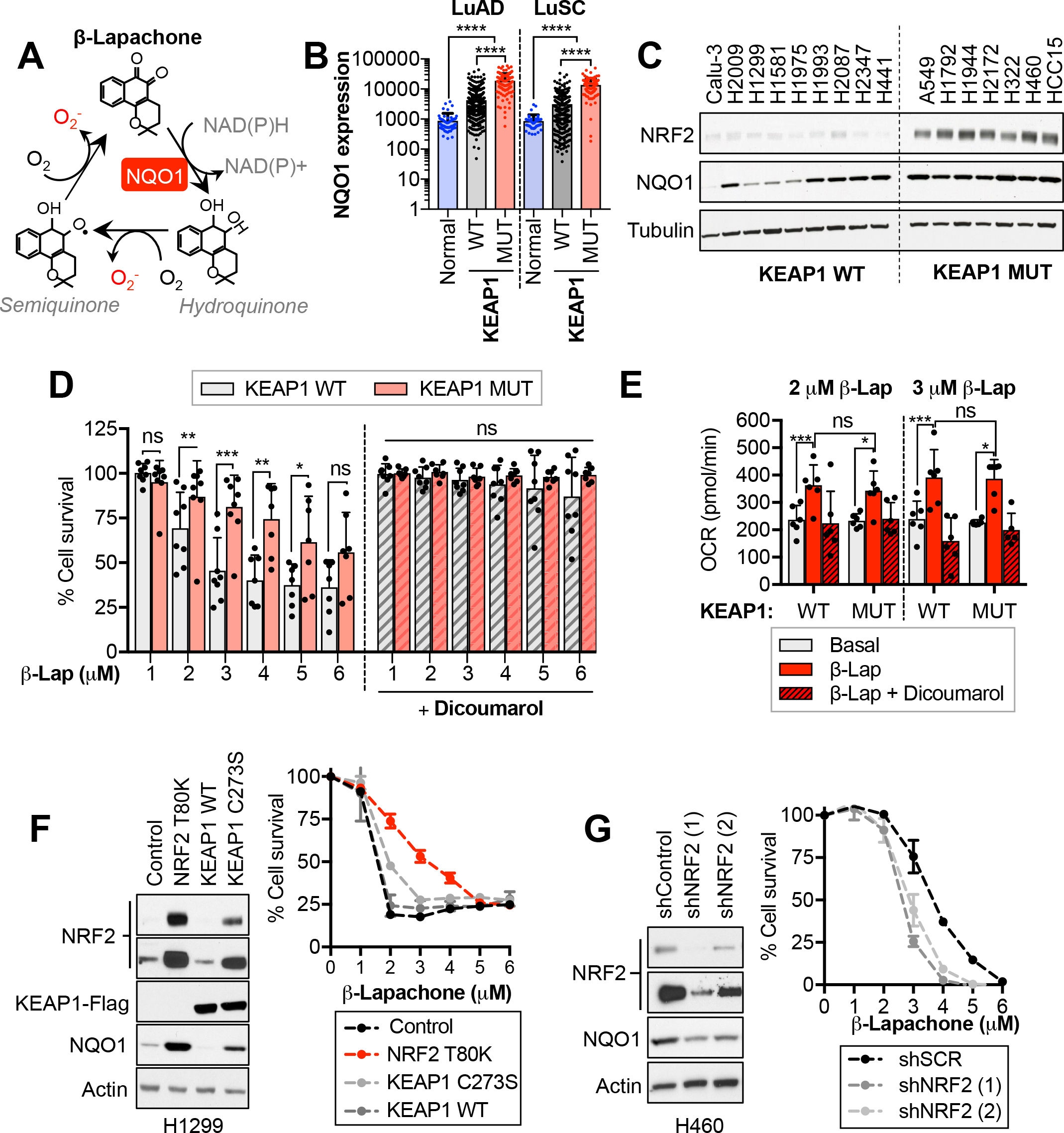
Aberrant activation of NRF2 increases resistance to β-Lapachone treatment. **(A)** Schematic representation of β-lapachone redox cycling. NQO1 catalyzes the two-electron reduction of β-lapachone to a hydroquinone form, which can spontaneously reoxidize, leading to the formation of superoxide radicals. **(B)** NQO1 mRNA expression in healthy lung tissue, lung adenocarcinomas (LuAD) and lung squamous cell carcinoma (LuSC). NQO1 mRNA expression in tumors was subdivided according to the KEAP1/NRF2 mutational status. **(C)** Western blot analyses of NRF2, NQO1 and Tubulin expression in a panel of wild-type (WT) and mutant (MUT) KEAP1 NSCLC cells. Note that Calu-3 cells harbor a polymorphic variant of NQO1 (NQO1*3, 465C<T). **(D)** Survival assays of KEAP1 wild-type and KEAP1 mutant (MUT) NSCLC cell lines exposed to β-Lapachone alone (left) or in combination with the NQO1 inhibitor dicoumarol (right). Cells were treated with the indicated concentrations of β-lapachone alone or in combination with dicoumarol (50 µM). **(E)** Oxygen consumption rates of a panel of KEAP1 WT and MUT NSCLC cells exposed to 2 and 3 µM of β-lapachone alone or in combination with 50 µM of dicoumarol for 117 minutes. **(F)** H1299 cells (KEAP1 WT) were infected with an empty vector (Control) or a virus coding for the expression of NRF2 T80K, KEAP1 wild-type or KEAP1 C273S. Left, western blot analyses of NRF2, Flag (for KEAP1 detection), NQO1 and actin (loading control). Right, survival assays of cells exposed to β-lapachone. **(G)** Western blot (left) and survival assays (right) of H460 (KEAP1 MUT) infected with virus coding for shRNAs against NRF2 (shNRF2) or an shRNA scramble control. *\*Note, for survival assays, cells were exposed to β-Lapachone for 2 hours, after which medium was replaced and cell viability was assessed 48 hours after treatment CellTiter-Glo (D) or crystal violet staining (F,G)*.

## Results

### Aberrant activation of NRF2 in NSCLC promotes resistance to B-lapachone

We compared the mRNA levels of NQO1 in healthy lung tissue, adenocarcinoma (LuAD) and squamous cell carcinoma (LuSC) patients using the TGCA dataset (Figure 1B). NQO1 expression in lung tumors was further subdivided according to the NRF2/KEAP1 mutational state. Both LuAD and SCC tumors had significantly higher NQO1 levels compared to normal lung tissue. Further, tumors harboring NRF2/KEAP1 mutations displayed significantly higher expression of NQO1 than those lacking mutations in this pathway. Remarkably, a number of KEAP1^WT^ NSCLC tumors exhibited elevated NQO1 levels, suggesting that NQO1 overexpression in KEAP1/NRF2^WT^ tumors can also result from alternative mechanisms of NRF2 activation (i.e. epigenetic silencing of KEAP1^27, 28^, oncogenic activation of NRF2^29^) or through NRF2-independent mechanisms.

To examine the influence of KEAP1/NRF2 mutations on NSCLC response to β-lapachone treatment, we assessed the cytotoxic efficacy of β-lapachone in a panel of sixteen NSCLC cell lines, seven of which harbor characterized inactivating mutations of KEAP1 (Figure 1C, S1A). In line with the mRNA data of LuSC and LuAD patients, KEAP1^MUT^ cell lines displayed uniformly high NQO1 protein levels, while protein levels of NQO1 in KEAP1^WT^ cells were highly variable. To determine the range of doses of β-lapachone that promote cell death in a NQO1-dependent manner, cells were treated with increasing concentrations of β-lapachone alone or in combination with the NQO1 inhibitor dicoumarol^30, 31^ (Figure 1D). Additionally, we included in the study Calu-3 cells, which harbor a polymorphic variant of NQO1 (NQO1*3) that results in 95% lower enzyme levels^32–34^ (Figure S1B). To recapitulate β-lapachone *in vivo* half-life conditions^35^, cells were treated with β-lapachone for two hours and cell viability was analyzed forty-eight hours after treatment. Our results showed that doses ranging from 1-6 µM induced cell death in a dose-dependent and NQO1-specific manner. Remarkably, KEAP1 mutation conferred resistance to β-lapachone treatment (Figure 1D).

To test the ability of KEAP1^WT^ and KEAP1^MUT^ cell lines to promote redox cycling of β-lapachone, we monitored the oxygen consumption rate (OCR) using the seahorse bioanalyzer. Basal OCR was monitored prior injection of β-lapachone, and OCR was followed for 2 hours after β-lapachone addition. To validate whether changes in the OCR were a consequence of NQO1-dependent redox cycling of β-lapachone, we also monitored the OCR of cells co-treated with β-lapachone and dicoumarol. Treatment with β-lapachone resulted in a significant increase of the OCR in both KEAP1^WT^ and KEAP1^MUT^ cells, which was precluded by the addition of dicoumarol (Figure 1E). Further, there was no significant difference in the OCR between KEAP1^WT^ and KEAP1^MUT^ cells after β-lapachone treatment, indicating that protein levels of NQO1 in KEAP1^WT^ NSCLC cells are not limiting for redox cycling of β-lapachone. Hence, both KEAP1^WT^ and KEAP1^MUT^ cancer cells are capable of NQO1-dependent redox cycling of β-lapachone at comparable rates.

To determine whether β-lapachone resistance was NRF2-dependent, H1299 cells (KEAP1^WT^) were infected with virus encoding for an inactivating mutation of KEAP1 (C273S) or a gain-of function NRF2 mutation (T80K), to promote the aberrant activation of NRF2^7, 36, 37^. In agreement with our previous findings, overexpression of KEAP1^C273S^ and NRF2^T80K^ but not KEAP1^WT^ led to the accumulation of NRF2, which promoted resistance to β-lapachone exposure (Figure 1F). Consistently, NRF2 silencing by shRNAs markedly reduced resistance to β-lapachone in H460 cells (KEAP1^MUT^) (Figure 1G). Additionally, we compared the β-lapachone sensitivity of NRF2-knockout A549 cells^38^ infected with a control vector or with virus coding for NRF2 expression (Figure S1C, S1D). In agreement with our previous data, NRF2-knockout cells exhibited increased sensitivity to β-lapachone compared to cells reconstituted for NRF2 expression. Collectively, these results indicate that aberrant activation of NRF2 in KEAP1 mutant lung cancer cells confers resistance to β-lapachone exposure.

### Activation of NRF2 promotes active scavenging of β-lapachone-induced ROS and attenuates DNA damage

Given the established role of NRF2 in protection against ROS through the transcriptional regulation of antioxidant enzymes, we evaluated whether KEAP1^MUT^ cells harbor an increased capacity to detoxify β-lapachone-induced ROS. We monitored ROS generation after β-lapachone exposure in our panel of NSCLC cell lines using the fluorogenic probe CellROX Green, as previously described^39^ (Figure 2A). We found that KEAP1^WT^ cells displayed a significantly higher fold-induction of ROS after 1-hour β-lapachone treatment compared to KEAP1^MUT^. To confirm whether β-lapachone promotes cell death via induction of ROS, we supplemented the media with exogenous catalase to increase the antioxidant capacity of the cells (Figure 2B). Consistent with prior studies^22, 40^, addition of exogenous catalase abrogated β-lapachone-induced cell death. Next, we assessed whether KEAP1^MUT^ cells were protected against β-lapachone-induced DNA damage^32, 39^. We monitored levels of phosphorylated H2A.X (γ-H2AX), a sensitive molecular marker of DNA damage. We observed a time-dependent accumulation of DNA damage in H1299 cells (KEAP1^WT^), while A549 cells (KEAP1^MUT^) did not exhibit increased γ-H2AX following 2-hour treatment with 3 µM β-lapachone (Figure 2C). We extended these analyses to a larger panel of cell lines, and we found that KEAP1^WT^ cells, but not KEAP1^MUT^ cells, accumulated γ-H2AX following β-lapachone exposure (Figure 2D). Accordingly, ectopic expression of KEAP1^C273S^ or NRF2T^80K^ in H1299 cells also promoted resistance to β-lapachone-induced DNA damage and decreased accumulation of ROS (Figure 2E,F and S2A).

**Figure 2.**
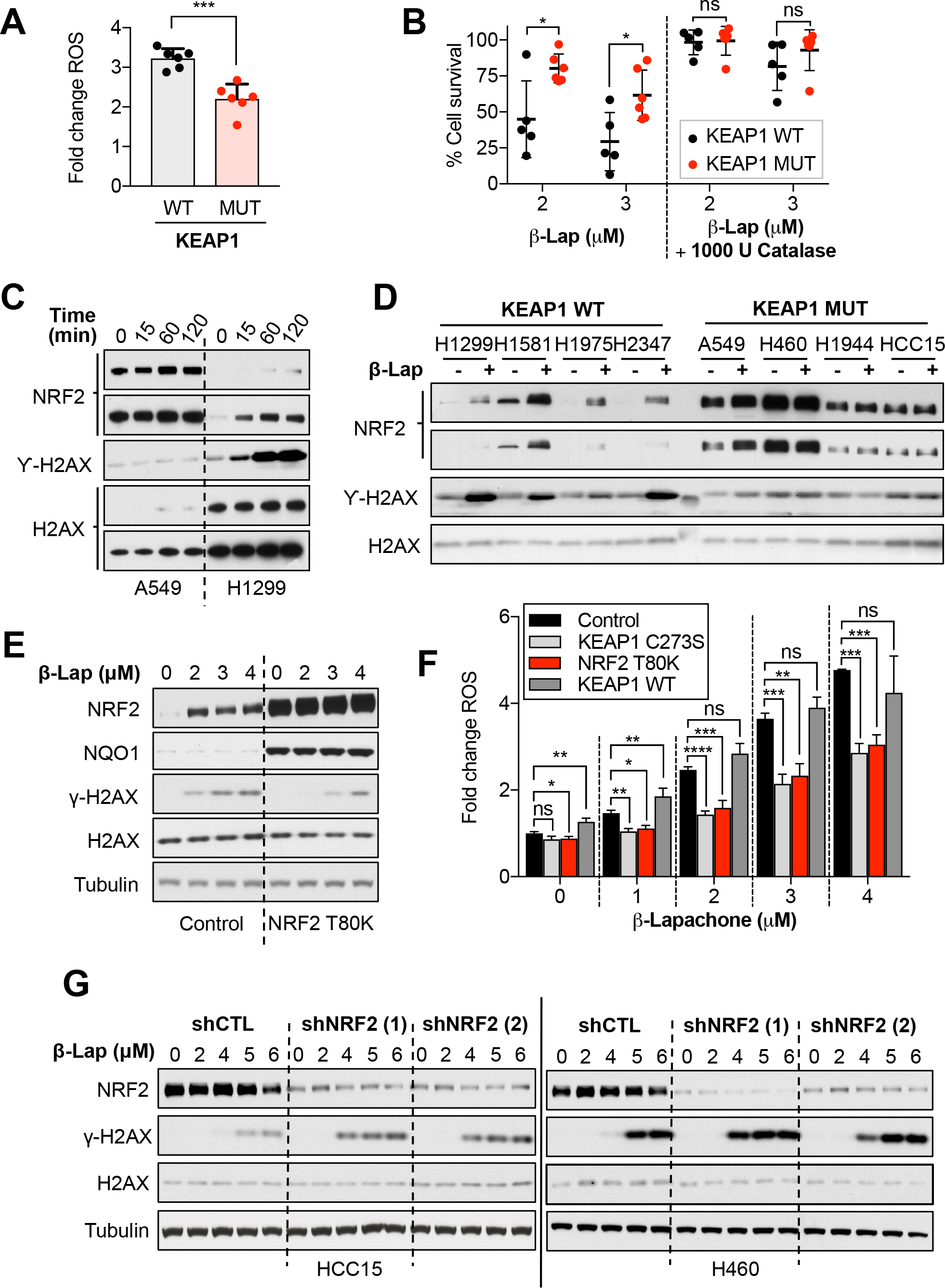
Activation of NRF2 promotes active scavenging of β-lapachone-induced ROS and attenuates DNA damage. **(A)** A panel of KEAP1 mutant and wild-type KEAP1 NSCLC cell lines were exposed to 3 µM of β-lapachone for one hour. Relative induction of ROS was measured using the green fluorescent probe CellROX green by flow cytometry. **(B)** Cell survival analyses of NSCLC cell lines exposed to β-lapachone alone or co-treated with 1000 U catalase/ well (96-well plate) for 2 hours. Surviving cells were stained with crystal violet 48 hours after treatment. **(C)** A549 and H1299 cells, KEAP1 mutant and wild-type respectively, were exposed to 3 µM of β-lapachone for 0, 15, 60 and 120 minutes, after which cells were collected and protein levels of NRF2, total H2AX (loading control), and the DNA damage marker (γ-H2AX, pS139) were assessed by western blotting. **(D)** Western blot analyses of nuclear extracts of a panel of KEAP1 wild-type (WT) or mutant (MUT) NSCLC cell lines. Cells were treated with 3 µM of β-Lapachone for 2 hours. Protein levels of NRF2, total H2AX (loading control), and the DNA damage marker (γ-H2AX, pS139) were assessed by western blotting. **(E)** DNA damage assessment of H1299 cells (KEAP1 WT) were infected with an empty vector (Control) or a virus coding for the expression of NRF2 T80K. Left, Cells were exposed to 2, 3 or 4 µM of β-lapachone. Protein levels of NRF2, NQO1, total H2AX (loading control), Tubulin (loading control) and the DNA damage marker (γ-H2AX, pS139) were assessed by western blotting. See supplementary figure 2A. **(F)** H1299 cells (KEAP1 WT) were infected with an empty vector (Control) or a virus coding for the expression of NRF2 T80K, KEAP1 wild-type or KEAP1 C273S. Cells were exposed to 1-4 µM of β-Lapachone for 1 hour and resulting fluorescent signal of the probe CellROX green was measured by flow cytometry. **(G)** H460 and HCC15 cells (KEAP1 MUT) infected with virus coding for shRNAs against NRF2 (shNRF2) or an shRNA scramble control were treated with the indicated concentration of β-lapachone for 2 hours, after which protein levels of NRF2, total H2AX (loading control), Tubulin (loading control) and the DNA damage marker (γ-H2AX, pS139) were assessed by western blotting.

We next assessed whether depletion of NRF2 could exacerbate β-lapachone-induced DNA damage in KEAP1^MUT^ cells (Figure 2G). Indeed, shRNA-mediated silencing of NRF2 in H460 and HCC15 cells increased γ-H2AX levels following β-lapachone exposure. Lastly, we interrogated ROS and DNA damage levels in NRF2-KO A549s infected with NRF2^WT^ or an empty control vector after β-lapachone treatment (Figure S2B, S2C). Consistently, NRF2-knockout cells exhibited greater production of ROS and accumulation of DNA damage markers compared to cells expressing NRF2. Together, these results demonstrate that constitutive activation of NRF2 in NSCLC protects cells from β-lapachone exposure by decreasing ROS-mediated DNA damage.

### Inhibition of thioredoxin-dependent systems but not catalase and glutathione overcome NRF2-mediated resistance to β-lapachone

To sensitize KEAP1^MUT^ lung cancer cells to β-lapachone treatment, we sought to identify and inhibit key NRF2-regulated antioxidant pathways. Given the major role of hydrogen peroxide in mediating β-lapachone toxicity, we tested whether inhibition of individual peroxide detoxification systems could overcome β-lapachone resistance (Figure S3A). First, we tested the relevance of catalase in the sensitivity to β-lapachone by using shRNAs (Figure 3A, 3B). We observed that depletion of catalase did not affect the β-lapachone sensitivity of NSCLC cells, regardless of KEAP1 mutational status. Of note, we found that H460 cells did not express detectable catalase protein (Figure 3B, S3B).

**Figure 3.**
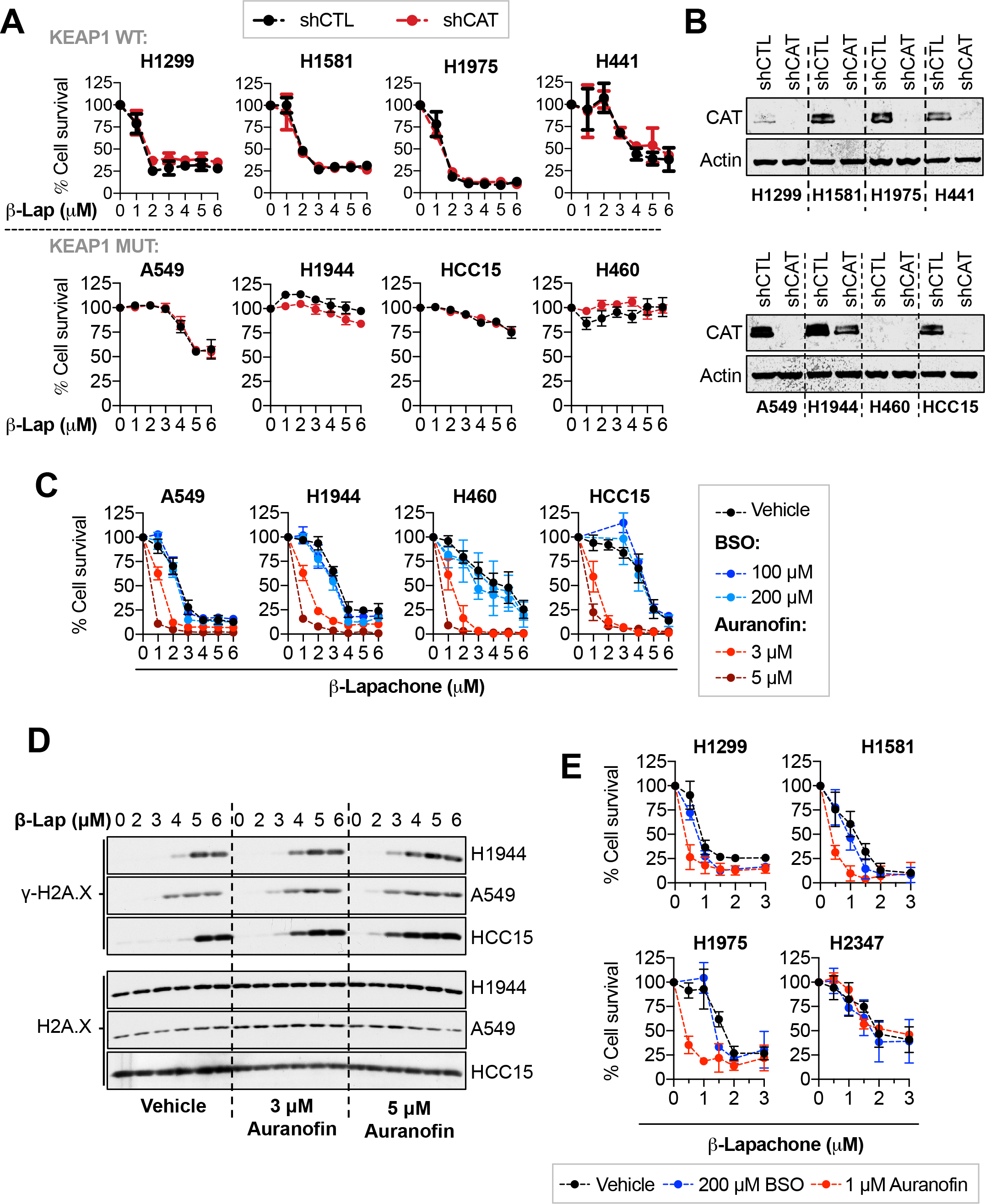
Inhibition of the TXN-dependent system but not GSH and catalase enhances sensitivity to B-lapachone treatment. **(A)** Assessment of β-lapachone sensitivity of NSCLC cell lines infected with shRNA against catalase (shCAT) or non-targeting control shRNA (shCTL). **(B)** Western blotting analysis of CAT and Actin (loading control) to validate the efficacy of the shRNAs against catalase. **(C)** Survival assays of a panel of KEAP1^MUT^ cells were exposed to β-lapachone alone or in combination with BSO (24 hours pre-treatment) or auranofin (2 hours co-treatment) to specifically inhibit the glutathione, and thioredoxin-dependent systems, respectively. **(D)** DNA damage assessment of a panel of KEAP1 mutant cells exposed to β-lapachone alone or in combination with 3 or 5 µM of auranofin. Protein levels of total H2AX (loading control) and the DNA damage marker (γ-H2AX, pS139) were assessed by western blotting. **(E)** Survival assays of KEAP1^WT^ cells exposed to β-lapachone alone or in combination with BSO (24 hours pre-treatment) or auranofin (2 hours co-treatment).

NRF2 is a major upstream transcriptional regulator of enzymes involved in the thioredoxin- and glutathione-dependent antioxidant systems, which share certain redundancy in detoxifying hydrogen peroxide. We tested whether individual inhibition of the thioredoxin (TXN)- or glutathione-dependent systems could overcome NRF2-mediated resistance to β-lapachone. To inhibit glutathione synthesis, we used buthionine sulfoximine (BSO), a well-characterized inhibitor of glutamate-cysteine ligase (GCL)^41^. Twenty-four-hour treatment with 100 or 200 µM of BSO depleted >95% of the total pool of glutathione (Figure S3C). To inhibit the TXN-dependent system, cells were treated with 1-5 µM auranofin, a pan-thioredoxin reductase (TXNRD) inhibitor for 2 hours^42^. We observed a dose-dependent inhibition of both the cytosolic and mitochondrial TXN-dependent systems in KEAP1^WT^ cells, as seen by oxidation of peroxiredoxin 1 and 3 (Figure S3D). Interestingly, KEAP1^MUT^ cell peroxiredoxins were resistant to oxidation, and KEAP1^WT^ NSCLC cells are more sensitive to auranofin-induced cell death in the absence of exogenous oxidants (Figure S3E). We next examined the effect of these inhibitors on β-lapachone sensitivity. Depletion of glutathione did not increase the sensitivity to β-lapachone treatment, while auranofin significantly increased β-lapachone-induced cell death and DNA damage in KEAP1^MUT^cells (Figure 3C **and 3D**). Similarly, auranofin but not BSO increased sensitivity of KEAP1^WT^ cells to β-lapachone exposure (Figure 3E).

These data suggest that although inhibition of the TXN-dependent system increases sensitivity of KEAP1^MUT^ cells to β-lapachone, the inherent reinforcement of this antioxidant pathway in KEAP1^MUT^ cells might represent a challenge *in vivo*. Further, single inhibition of the glutathione or catalase it is not sufficient to increase sensitivity to β-lapachone treatment, suggesting that inhibition of these pathways is compensated by other antioxidant mechanisms.

### Inhibition of SOD1 potentiates β-lapachone anti-tumor efficacy in KEAP1/NRF2^MUT^ NSCLC

The inherent redundancy in the peroxide detoxification systems represents a major challenge to sensitize KEAP1^MUT^ cells to ROS generators, as inhibition of one of these pathways may be compensated by other antioxidant enzymes. In contrast, SOD1 has a unique role in catalyzing the dismutation of cytosolic superoxide radicals generated by β-lapachone (Figure 4A). Given this unique role of SOD1, we interrogated the effects of SOD1 inhibition on β-lapachone efficacy. We infected KEAP1^MUT^ cells with virus coding for shRNAs against SOD1 or a non-targeting shRNA (Figure S4A). Depletion of SOD1 markedly increased β-lapachone-mediated cell death in KEAP1^MUT^ cell lines and increased DNA damage (Figure 4B, 4C, S4B). Importantly, β-lapachone treatment did not change SOD1 activity (Figure S4C).

**Figure 4.**
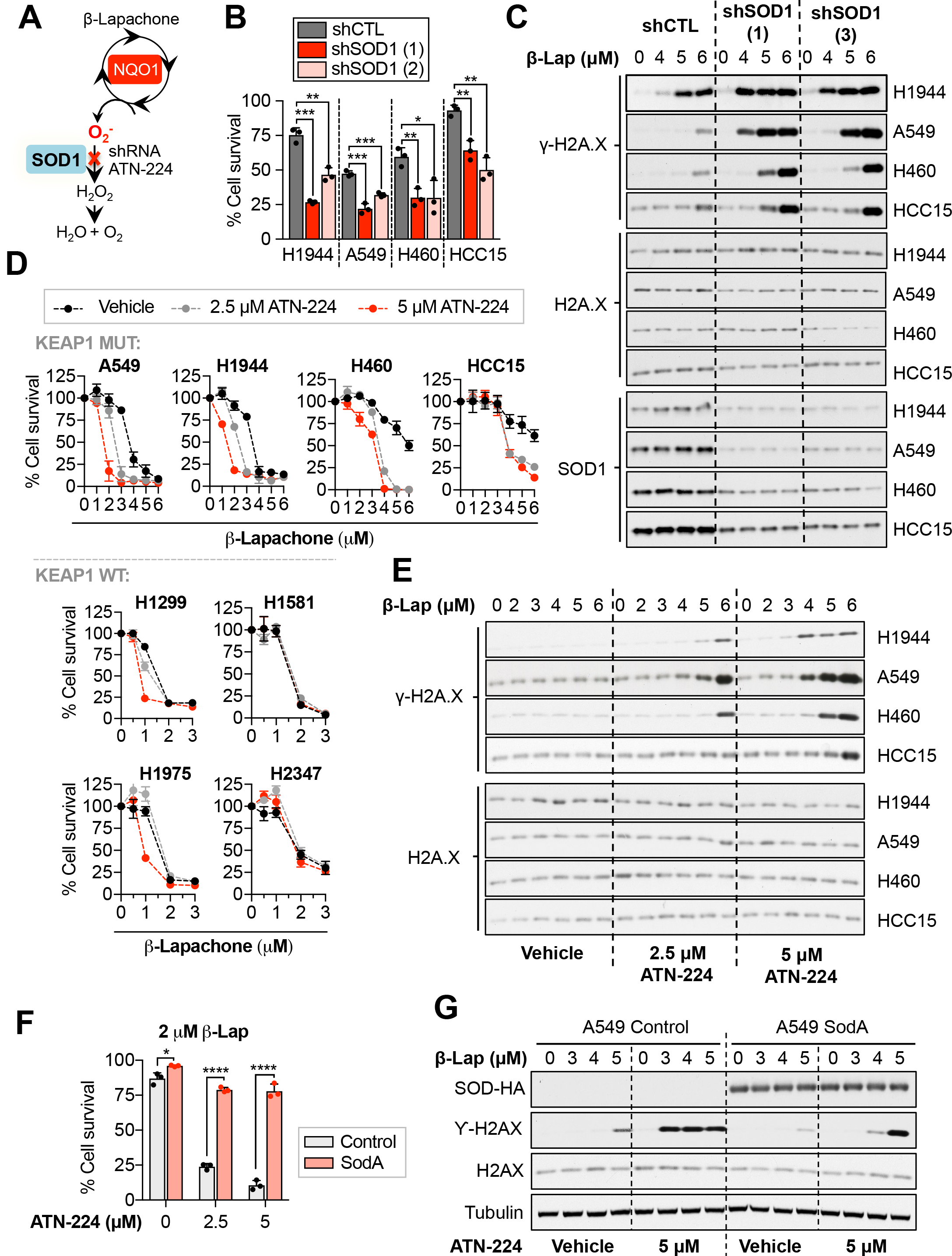
Inhibition of the copper-zinc superoxide dismutase (SOD1) potentiates β-lapachone anti-tumor efficacy in NSCLC *in vitro*. **(A)** Schematic representation of the potential role of SOD1 in the detoxification of β-lapachone-induced ROS. **(B)** Survival assays of a panel of KEAP1 mutant NSCLC cells infected with virus encoding for shRNAs against SOD1 (1, 2) or with an empty vector (CTL) as control. NSCLC cells were treated with vehicle (0.012% DMSO) or with 3 µM β-lapachone for 2 hours. Cell viability was assessed 48 hours after treatment. **(C)** KEAP1 mutant cells were infected with virus encoding for shRNAs against SOD1 (1, 3) or with an empty vector (CTL) as control. Cells were treated with 3 µM β-lapachone and protein levels of total H2AX (loading control), SOD1 and the DNA damage marker (γ-H2AX, pS139) were assessed by western blotting 2 hours after treatment. **(D)** A panel of KEAP1 mutant (top) and KEAP1 WT (bottom) NSCLC cells were treated for 24 hours with vehicle (0.05% DMSO) or with 2.5 or 5 µM of ATN-224, after which cells were treated with β-lapachone for 2 hours in ATN-224 containing medium. Fresh media was added after treatment to allow for SOD1 reactivation. Cell viability was assessed 48 hours after treatment using crystal violet. **(E)** NSCLC cells were treated for 24 hours with vehicle (0.05% DMSO) or with 2.5 or 5 µM of ATN-224, after which cells were treated with β-lapachone for 2 hours in ATN-224 containing medium. Protein levels of total H2AX (loading control) and the DNA damage marker (γ-H2AX, pS139) were assessed by western blotting. **(F)** A549 cells were infected with virus coding for the expression of E. coli MnSOD (SodA) or an empty vector (control). Cells were treated with β-lapachone alone or in combination with the copper chelator ATN-224. Surviving cells were stained with crystal violet 48 hours after treatment. **(G)** A549 SodA or control were pre-treated for 24 hours with ATN-224 (5 µM) or vehicle, after which cells were exposed to the indicated concentrations of β-lapachone for 2 hours. Protein levels of SodA (HA-tag), Tubulin (loading control), total H2AX (loading control), and the DNA damage marker (γ-H2AX, pS139) were assessed by western blotting.

To validate these findings, we sough to test the effect of pharmacological inhibition of SOD1 on β-lapachone efficacy. However, direct inhibitors of SOD1 that have been shown efficacy in cell-based assays or *in vivo* are lacking. Importantly, the cytosolic (SOD1) and the extracellular (SOD3) superoxide dismutases require copper and zinc to function, while the mitochondrial isoform (SOD2) relies on manganese. Thus, indirect inhibition of SOD1 can be achieved through copper chelation^43, 44^, while the mitochondrial SOD activity should remain intact.

First, we validated whether we could achieve SOD1 inhibition in cell culture using the copper chelator ATN-224 (Figure S4D). We observed that most of SOD1 activity was inhibited after twenty-four-hour treatment using 2.5-5 µM. As expected, copper chelation did not affect SOD2 activity. Inhibition of SOD1 resulted in increased sensitivity of KEAP1^MUT^ cells to β-lapachone-mediated cell death and DNA damage (Figure 4D **and 4E**). Of note, SOD1 inhibition had little to no effect in KEAP1^WT^ cells (Figure 4D). We also examined the protein levels and activity of SOD1 across our panel of NSCLC cells to evaluate whether KEAP1^MUT^ cells have higher capacity to detoxify cytosolic superoxide (Figure S4E). KEAP1 mutant cells did not exhibit significantly higher levels of SOD1, suggesting that inactivation of KEAP1 does not confer a reinforced capacity to detoxify cytosolic superoxide radicals through SOD1 upregulation.

To confirm whether copper chelation sensitizes KEAP1^MUT^ cells through SOD1 inhibition, A549 cells were engineered to express an *Escherichia coli* manganese-dependent superoxide dismutase enzyme (SodA) to rescue SOD activity. Importantly, SodA expression significantly rescued the effects of copper chelation on β-lapachone treatment (Figure 4F and 4G). Altogether, these data demonstrate that inhibition of SOD1 selectively increases β-lapachone efficacy in KEAP1^MUT^ NSCLC cells.

## Discussion

Aberrant NRF2 activation promotes resistance to therapeutics that rely on the production of ROS, including multiple chemotherapeutics and radiation therapy. In this study, we find that NRF2 activation also promotes resistance to the NQO1-activatable prodrug β-lapachone, which relies on the generation of superoxide for its efficacy. While direct NRF2 inhibition could potentially reverse this resistance, NRF2 inhibitors identified to date either lack specificity or potency. Further, the effects of NRF2 whole-body inhibition as anti-tumor strategy remain unclear, as NRF2 activity in necessary for normal functioning of immune cells^45, 46^.

Consequently, we have evaluated whether inhibition of the cellular antioxidant systems can reverse the resistance of NRF2 active cells to ROS. We find that inhibitors of the TXN-dependent peroxide detoxification system and SOD1, but not glutathione or catalase depletion, can reverse the resistance of KEAP1^MUT^ cells to β-lapachone. Surprisingly, we find that KEAP1^MUT^ cells were highly resistant to auranofin compared to KEAP1^WT^ cells, which raises a concern about the toxicity of the required auranofin doses to healthy tissues. The resistance of KEAP1^MUT^ cells is likely explained by the redundancy of the thioredoxin- and glutathione-dependent antioxidant systems, which can both detoxify H_2_O_2_ and protect from excessive oxidation. Indeed, even at the highest doses of auranofin, KEAP1^MUT^ cells exhibited no peroxiredoxin oxidation in the absence of exogenous oxidants. This redundancy of the antioxidant program, particularly in cancer cells, represents a major challenge^47^. Although concomitant inhibition of the GSH and the TXN-dependent systems has been shown to abolish such redundancy *in vitro*, this strategy appears to be highly toxic and potentially lethal *in vivo*. Elias Arnér’s laboratory recently previously reported that combination of auranofin and BSO in mice, at concentrations that are tolerable as single agents, was lethal after the first round of administration^48^.

While previous studies demonstrated an important role for H_2_O_2_ in mediating β-lapachone cytotoxicity, we uncovered a surprising role for SOD1 in the protection against cell death in KEAP1^MUT^ cells. The selectivity of SOD1 inhibition for KEAP1^MUT^ cells may be a consequence of the enhanced peroxide detoxification capacity of these cells, while KEAP1^WT^ cells may be equally sensitive to both peroxide and superoxide. Importantly, our results suggest that antioxidant inhibition may sensitize NRF2/KEAP1 mutant cells to other ROS-generating therapeutics to which these cells are generally resistant. Importantly, a recent study found that SOD1 deletion or modulation of copper availability sensitized Jurkat cells to the superoxide generating compound paraquat^49^. Interestingly, SOD1 inhibition alone could target NSCLC cells in a study using A549 and H460 cells, which are KEAP1 mutant^50^. This study also found SOD1 inhibition was synergistic with glutathione depletion, suggesting that alternative combinations to inhibit the antioxidant capacity of KEAP1 mutant cells may be possible.

Auranofin and ATN-224 join the growing list of agents that can sensitize cells to β-lapachone. These agents converge on DNA damage and repair (PARP and XRCC1)^32, 39^, NAD+ availability (NAMPT)^51^, and ROS (GLS1 inhibition)^52^. Recently, more potent NQO1-activatable compounds, including the deoxynyboquinones (DNQs) have shown equivalent efficacy to β-lapachone at 6-fold lower dose^22^. Isobutyl-deoxynyboquinone (IB-DNQ) is being developed for clinical use^53^, and lacks the issues with stability, solubility and red blood cell toxicity that β-lapachone demonstrates. Our results suggest KEAP1^MUT^ cells would be resistant to IB-DNQ as well, and KEAP1/NRF2 mutation status should be considered in addition to NQO1 levels when predicting *in vivo* response to these agents.

## Materials and Methods

### Reagents and chemicals

β-Lapachone and dicoumarol were a gift from Professor David Boothman. Nitrotetrazolium Blue chloride powder (N6639), riboflavin (R7649), Auranofin (A6733-10MG) and Catalase from bovine liver (C1345) were all obtained from Sigma Aldrich. L-Buthionine-(S,R)-Sulfoximine (BSO) was obtained from Cayman Chemical (14484) or from Sigma Aldrich (B2515-500MG). ATN-224 was purchased from Cayman Chemical (23553). N-Ethylmaleimide was purchased from Chemimpex.

### Cell culture and reagents

Parental NSCLC cell lines were previously described (DeNicola et al., 2015). NRF2-KO A549 cells were obtained from Dr. Laureano de la Vega (Torrente et al., 2017). Cell lines were routinely tested and verified to be free of mycoplasma (MycoAlert Assay, Lonza). All lines were maintained in RPMI 1640 media (Hyclone or Gibco) supplemented with 5-10% FBS without antibiotics at 37°C in a humidified atmosphere containing 5% CO_2_ and 95% air. Lenti-X 293T cells were obtained from Clontech, and maintained in DMEM media (Hyclone or Gibco) supplemented with 10% FBS.

### Antibodies

The following antibodies were used: NRF2 (Cell Signaling Technologies, D1Z9C, Cat #12721), NQO1 (Sigma Aldrich, Cat# HPA007308), β-actin (Thermo Fisher, clone AC-15, Cat # A5441), α-tubulin (Santa Cruz, TU-02, Cat #sc-8035), Total Histone H2A.X (Cell Signaling Technologies, D17A3, Cat #9718), Gamma Histone H2A.X (Cell Signaling Technologies, 20E3, Cat #7631S), KEAP1 (Millipore Sigma, Cat# MABS514), SOD1 (Cell Signaling Technologies, 71G8, Cat #4266), Catalase (Cell Signaling Technologies, D4P7B, Cat #12980S), Flag Tag (Cell Signaling Technologies, M2, Cat# 14793S), HA-Tag (Cell Signaling Technologies, C29F4, Cat #3724), Prdx3 (Abcam, Cat #ab73349), Prdx1 (Cell Signalling, D5G12, Cat# 50-191-580)

### Plasmids

shRNAs against SOD1, NRF2 and catalase in the pLKO.1 backbone were purchased from Dharmacon. shSOD1(1) TRCN0000039809, shSOD1 (2) TRCN0000039812, shSOD1 (3) TRCN0000039808. shNRF2(1) TRCN0000007555 and shNRF2 (2) TRCN0000281950 were previously described^54^. shCAT TRCN0000061756. The shRNA control was purchased from Sigma-Aldrich (#SHC002). pLX317-NRF2 and pLX317-NRF2^T80K^ were obtained from Dr. Alice Berger^55^ and the pLX317 empty control vector was generated by site-directed mutagenesis as previously described^56^. SodA: The cDNA encoding for the MnSOD (Gene name: sodA) of Escherichia coli (strain K12) containing a c-terminal Influenza Hemagglutinin (HA) reporter tag was cloned into the backbone pLX317-empty using the In-Fusion cloning kit (Clontech). The backbone was cut with BamHI and EcoRI.

### Stable cell line generation

Stable cell lines were generated via lentiviral transduction. For lentivirus production, Lenti-X 293T cells (Clontech) were transfected at 90% confluence with JetPRIME (Polyplus) or Polyethylenimine (PEI). Packaging plasmids pCMV-dR8.2 dvpr (addgene # 8455) and pCMV-VSV-G (addgene #8454) were used. Cells were transduced for 6 hours by recombinant lentiviruses in growth media using polybrene (8 µg/ml). Twenty-four hours after transduction, puromycin (1 µg/ml) was added to the growth medium for 72 hours to select for infected cells.

### Cell viability assays

NSCLC cells were seeded in 96-well plates at a density of 2,500-5,000 cells/well in a 200 µl final volume. The following day, the media was replaced with 150 µl of fresh media containing the indicated concentrations of β-lapachone or vehicle (≤ 0.1% DMSO) for two hours, after which media was replaced. Two hours after treatment, medium was replaced by 200 µl fresh medium. Cell viability was assessed 48 hours after treatment with CellTiter-Glo (Promega) or crystal violet staining. To stain surviving cells with crystal violet, cells were washed in ice-cold PBS, fixed with 4% paraformaldehyde, stained with crystal violet solution (0.1% Crystal Violet, 20% methanol), washed with H_2_O and dried overnight. Crystal violet was solubilized in 10% acetic acid for 30 minutes and the OD600 was measured. Relative cell number was normalized to vehicle treated cells.

For experiments using the copper chelator ATN-224, cells were pre-treated with 1-5 µM of ATN-224 for 24 hours prior β-lapachone treatment. To ensure that no copper was added back with the media when cells were treated with β-lapachone, additional ATN-224 containing medium was prepared the day before β-lapachone treatment and stored at 4**°**C. After 24 hours, β-lapachone was prepared in ATN-224 containing medium and 150 µl were dispensed in each well. Similarly, cells were pre-treated for 24 hours with 100-200 µM of BSO prior β-lapachone exposure, which was prepared in BSO containing media to prevent the recovery of glutathione. Auranofin was added to the media in combination with β-lapachone for 2 hours.

### Protein extraction and Immunoblotting

Lysates were prepared in RIPA lysis buffer (20 mM Tris-HCl [pH 7.5], 150 mM NaCl, 1 mM EDTA, 1 mM EGTA, 1% NP-40, 1% sodium deoxycholate) containing protease and phosphatase inhibitors. To assess total/gamma-H2A.X levels in whole cell lysates, cells were lysed in boiling 1% w/v SDS RIPA lysis buffer supplemented with phosphatase and protease inhibitors. Cells were seeded in 6-well plates at 70-90% confluence. After drug treatment (β-lapachone +/-ATN-224 or auranofin) cells were washed in PBS and 300 µl of pre-warmed lysis buffer (90**°**C) was directly added to the wells. Cell lysates were transferred to a microcentrifuge tubes, incubated at 90**°**C for 5 minutes, followed by sonication to shred the DNA in a water bath sonicator (Diagenode). Samples were centrifuged at 13,000 x rpm for 15 minutes at 4**°**C to precipitate the insoluble fraction. The supernatant was transferred to a clean Eppendorf tube. Alternatively, total/gamma-H2A.X levels were monitored in nuclear extracts (see protocol below). Cell lysates were mixed with 6X sample buffer containing β-ME and separated by SDS-PAGE using NuPAGE 4-12% Bis-Tris gels (Invitrogen), followed by transfer to 0.45µm Nitrocellulose membranes (GE Healthcare). The membranes were blocked in 5% non-fat milk in TBS-T, followed by immunoblotting.

### Nuclear isolation

Cells were plated in 6-cm dishes at 70-90% confluence (5×10^5^ cells/well). The following day, cells were washed with ice-cold PBS, collected in 1 ml of ice-cold PBS, transferred to microcentrifuge tubes, and subjected to centrifugation at 13,000 × rpm for 1 min at 4**°**C. The cell pellet was resuspended in 400 µl of ice-cold of the low-salt buffer A (10 mM HEPES/KOH pH 7.9, 10 mM KCl, 0.1 mM EDTA, 0.1 mM EGTA and protease/phosphatase inhibitors). After incubation for 10 minutes on ice, 10 µl of 10% NP-40 was added and cells were lysed by gently vortexing. The homogenate was centrifuged for 10 seconds at 13,200 rpm. The supernatant was collected as the cytoplasmic fraction while the pellet containing the cell nuclei was washed 4 times in 400 µl buffer A. Cell nuclei was lysed in 100µl high-salt buffer B (20mM HEPES/KOH pH7.9, 400mM NaCl, 1mM EDTA, 1mM EGTA and protease/phosphatase inhibitors). The lysates were sonicated and centrifuged at 4°C for 15 minutes at 13,200 rpm. The supernatant representing the nuclear fraction was collected and protein concentration of both fractions was determined using the DC protein assay (Biorad). Cytosolic and nuclear fractions were further diluted to the desired protein concentration using the corresponding lysis buffers. One volume of 5X sample SDS loading buffer (250 mM Tris-Cl [pH 6.8], 10% [v/v] SDS, 40% [v/v] glycerol, and 0.1% [w/v] bromophenol blue, 15% [v/v] β-mercaptoethanol) was added to 4 volumes of the lysate and subjected to SDS-PAGE.

### In-gel SOD activity assay

Cells were plated in 35 mm diameter cell culture dishes at a density of 3.5 × 10^5^ if assayed the following day or 3.5 x 10^5^ cells/well if 24 hours of ATN-224 pre-treatment was required. Cells were collected by scraping with ice-cold PBS and lysed on ice for 30 min in 100 µl of SOD lysis buffer (8.96 mM Na_2_HPO_4_, 0.96 mM NaH_2_PO_4_, 0.1% Triton X-100, 5 mM EDTA, 5 mM EGTA, 50 mM NaCl, 10 % Glycerol). Samples were centrifuged for 15 minutes at 13,000 x rpm at 4°C, and supernatant transferred to a microcentrifuge tube. Protein concentration was determined using the DC protein Assay (Biorad). Protein samples were diluted to 0.7-1 µg/µl using the SOD lysis buffer and 1/5th volume of 5x native loading dye [0.31 M Tris pH 6.8, 0.05% bromophenol blue (w/v), 50% glycerol (v/v)]. Samples were subjected to native polyacrylamide gel electrophoresis (7.5%) at 90V for 2 hours at 4°C in a buffer containing 25 mM Tris, 192 mM glycine. The gel was stained in SOD staining solution (0.22 M K_2_HPO_4_, 0.02 M KH_2_PO_4_, 1% [w/v] Riboflavin, 1.3% [w/v] Nitro blue tetrazolium and 0.1% TEMED) in the dark for 1 hour, after which the gel was washed in dH_2_O and exposed to light until developed to dark blue/black (1 hour approximately). Achromatic bands correspond to SOD1 and SOD2.

### Reactive oxygen species measurement

Reactive oxygen species were measured using the CellROX green reagent (Thermofisher) according to the manufacturer’s instructions. In short, cells were plated in 24-well plates at 70% confluence. The following day, cells were treated with the indicated concentrations of β-lapachone in a 400 µl/well final volume for 30 minutes at 37°C. CellROX green reagent was added to a final concentration of 5 µM to the cells without replacing the media, and incubated for additional 30 minutes at 37°C. Cells were washed in PBS, trypsinized and transferred to microcentrifuge tubes. The cell suspension was centrifuged for 20 seconds at 13,000 x rpm, and resuspended in 300 µl of ice-cold PBS. The Green fluorescence of dye-loaded cells was determined by flow cytometry using the FACSCalibur flow cytometer (BD Biosciences) and the acquisition software CellQuest Pro. The mean fluorescence intensity of 10,000 discrete events was calculated for each sample.

### Glutathione measurement

H1944 and HCC15 cells were seeded on flat, round bottomed 96-well plates at a density of 1×10^4^ cells/well. The following day, media was replaced with media containing 100-200 µM of BSO or fresh media (negative control). The following day, the GSH-Glo™ Glutathione Assay (Promega, Cat# V6911) was used to measure the intracellular reduced glutathione pool (GSH). Data were normalized by percentage relative to the control non-treated sample.

### Redox western blotting

The protocol for determining the ratio of reduced and oxidized peroxiredoxin 1 and 3 was adapted from Prof. Mark Hampton’s lab^57^. One-day prior to the assay, cells were seeded in 6-well plates at 70-90% confluence (∼5×10^5^ cells/well). The following day, cell culture media was replaced with 1ml of media containing 1, 3 or 5 µM of auranofin or fresh medium. Final DMSO concentration was 0.013% (V/V). Two hours after treatment, media was aspirated and cells were gently washed with 1 ml of ice-cold PBS. 1 mg/ml of bovine catalase was added to the alkylation buffer (40mM HEPES, 50mM NaCl, 1mM EGTA, complete protease inhibitors, pH 7.4) 30 minutes prior collection. Immediately prior sample collection, 200 mM of N-Ethylmaleimide (NEM) were added to the alkylation buffer and the lysis buffer was warmed to 42°C for 1-2 minutes to dissolve the NEM. To lyse the cells, 200 µl of alkylation buffer were dispensed in each well, followed by 10 min incubation at room temperature. A solution of 10% CHAPS was added to the lysates to a final concentration of 1% CHAPS (20 µl/sample). Cell lysates were transferred to a 1.5 ml microcentrifuge tube, samples were vortex and incubated on ice for further 30 minutes, followed by 15 minutes centrifugation at 13,000 x rpm, 4**°**C. The supernatant was transferred to a clean 1.5 ml microcentrifuge tube. These redox western samples were mixed with a 4X non-reducing buffer prior to separation by SDS-Page.

### Analysis of NQO1 mRNA expression in patient samples

Patient normal lung, LuAD and LuSC data mRNA data was obtained by combining the data available in cBioportal and the MethHC databases. Patient LuAD and LuAD data from The Cancer Genome Atlas (TCGA), with associated KEAP1 mutation status (Illumina HM450 Beadchip), was obtained from cBioPortal^58, 59^. Patient IDs were matched with the data from The Cancer Genome Atlas (TCGA) via the MethHC database^60^, and expression of NQO1 in normal lung tissue data was included in the study.

### Seahorse assay (Oxygen consumption)

Seahorse assays were performed using the Seahorse XFe96 analyzer (Agilent) accoding to the manufacturer’s instructions. 4×10^4^ cells/well were seeded in the seahorse XF96 cell culture microplates at a final volume of 80 µl. Sensor cartridge was hydrated overnight in a non-CO_2_ 37°C incubator with Seahorse the XF calibrant (200 µl). The following day, cell media was changed to bicarbonate-free supplemented with 4.5 g/L glucose, 2mM glutamine and antibiotics (penicillin/streptomycin) at a final volume of 175 µl/well. Cells were incubated for 40 min-1 hour in a non-CO_2_ 37°C incubator. β-Lapachone and dicoumarol were prepared as concentrated stocks (8X). Three baseline measurements were obtained prior injection of β-Lapachone alone or in combination with dicoumarol. Non-treated cells were injected with vehicle (0.016% DMSO or 15 mM NaOH). After β-Lapachone/dicoumarol injections, the Oxygen Consumption Rate was followed for 2 hours (20 measurements).

### Statistical analyses

Data were analyzed using a two-sided unpaired Student’s t test. GraphPad Prism 7 software was used for all statistical analyses, and values of p < 0.05 were considered statistically significant (*P < 0.05; **P < 0.01; ***P < 0.001, **** P < 0.0001).

## Acknowledgements

We would like to express our sincere gratitude to Dr. Donita C. Brady and Julianne Davis for their thoughtful insights and technical advice. This work is supported by grants from the American Lung Association, NIH (R37-CA230042) and Florida Department of Health Bankhead-Coley research program (9BC07) to G.M.D. This work has also been supported by the Lung Cancer Center of Excellence and Flow Cytometry Core Facility at the Moffitt Cancer Center, an NCI designated Comprehensive Cancer Center (P30-CA076292).

## Author Contributions

Conceptualization, L.T. and G.M.D.; Methodology, L.T., and G.M.D.; Investigation, L.T., N.P., and A.F.; Resources – D.A.B; Writing – Original Draft, L.T. and G.M.D.; Writing – Review & Editing, L.T. N.P. and G.M.D.; Funding Acquisition, G.M.D. and E.B.H.; Supervision, G.M.D., E.B.H. and D.A.B.

## Declaration of interests

The authors declare no competing interests.

### Materials & Correspondence

Correspondence and material requests should be addressed to.

---

**Supplementary Figure 1.**
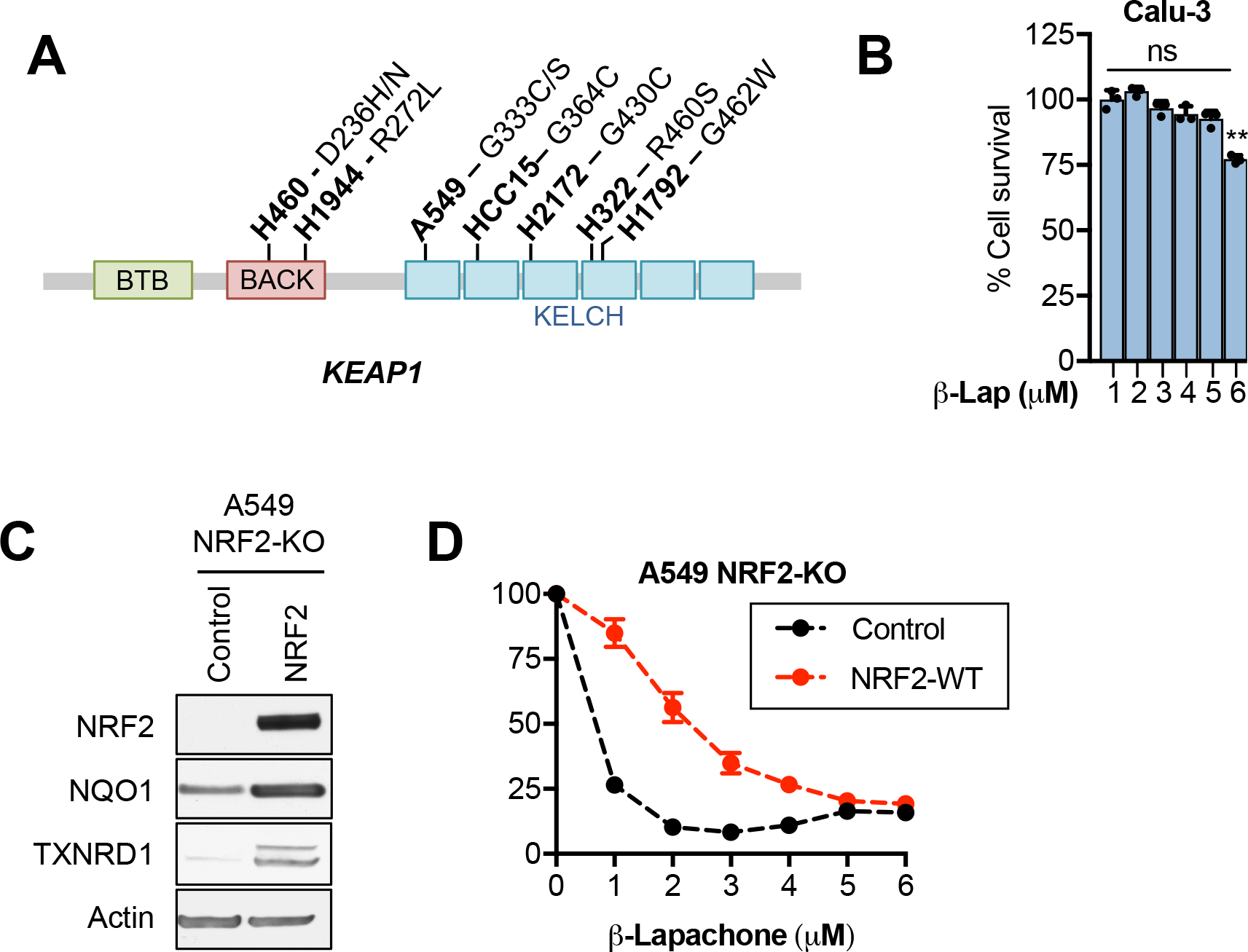
Alterations in the NRF2/KEAP1 pathway confer resistance to β-lapachone exposure. **(A)** Schematic representation of the KEAP1 mutations of the NSCLC cells included in our analyses. **(B)** Survival assays of Calu-3 (NQO1*3) exposed to β-lapachone for 2 hours. Cell viability was assessed 48 hours after treatment. **(C)** Western blot analyses of CRISPR-Cas9 engineered NRF2-KO cells infected with lentivirus coding for wild-type NRF2 or an empty vector (control). Cells were incubated with the puromycin selection agent for 72 hours. **(D)** Survival assays of A549 and H1299 NRF2-KO cells transduced with an empty vector (control) or an expression vector coding for NRF2 (NRF2) exposed to the indicated concentrations of β-lapachone for 2 hours, after which media was replaced and cell survival was analyzed 48 hours after treatment using CellTiter-Glo.

**Supplementary figure 2.**
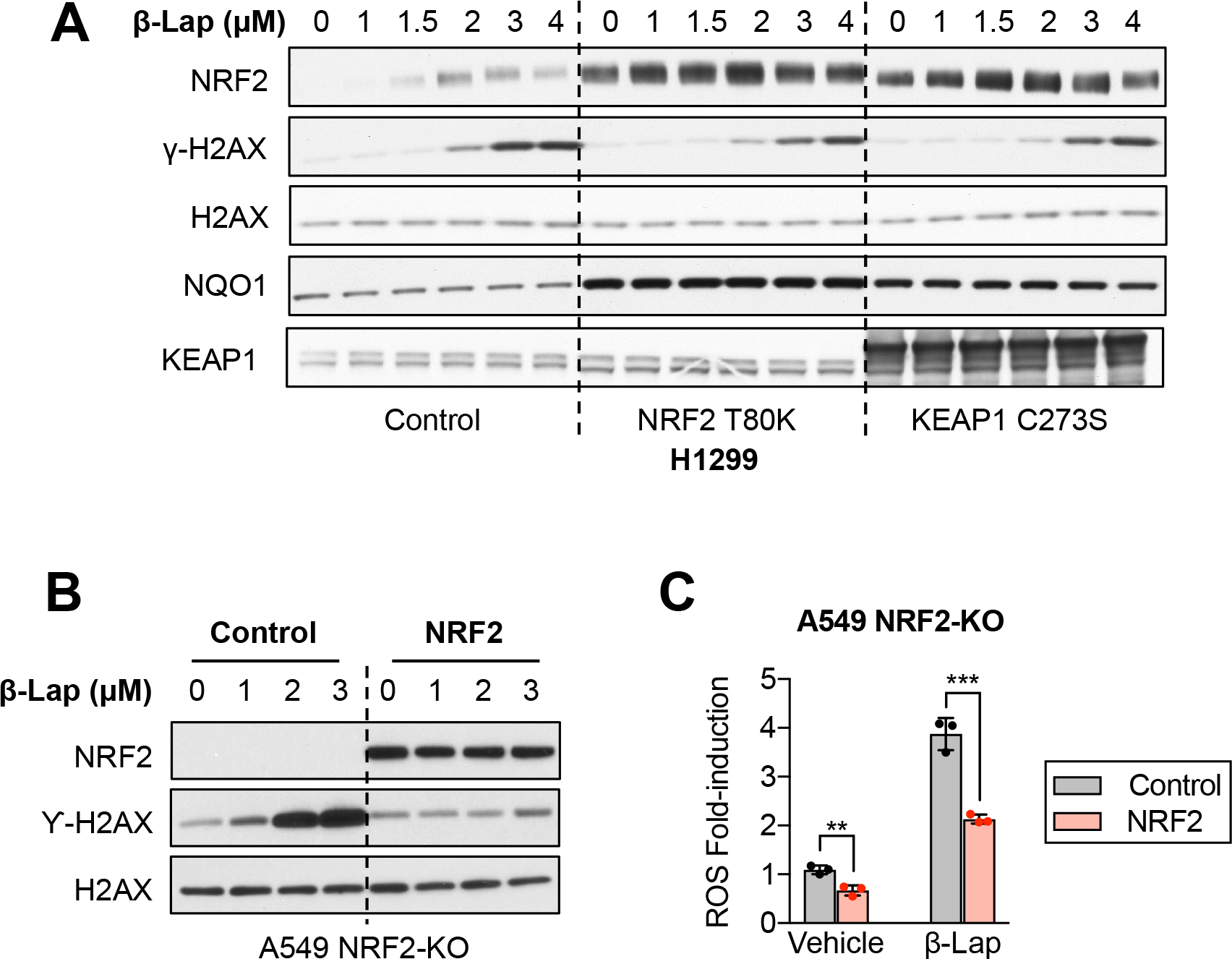
Activation of NRF2 renders NSCLC cells resistant to β-lapachone-induced DNA damage and promotes ROS scavenging. **(A)** H1299 cells (KEAP1 WT) were infected with virus coding for NRF2-T80K, KEAP1 C273S or an empty control as vehicle. Cells were exposed to the indicated concentrations of β-lapachone for 2 hours. Protein levels of NRF2, NQO1 KEAP1, total H2AX (loading control), tubulin (loading control) and the DNA damage marker (γ-H2AX) were assessed by western blotting. **(B)** DNA damage assessment in A549-KO cells control or reconstituted with NRF2 and exposed to β-lapachone for 2 hours. Protein levels of NRF2, total H2AX (loading control), and the DNA damage marker (γ-H2AX) were assessed by western blotting. **(C)** A549 NRF2-KO cells infected with an empty vector (control) or with a vector encoding for the expression of NRF2 were treated with 1 µM of of β-lapachone. ROS levels were measured using CellROX green and fluorescence was measured by flow cytometry. P-values: Vehicle= 0.006; β-Lapachone= 0.0009

**Supplementary figure 3.**
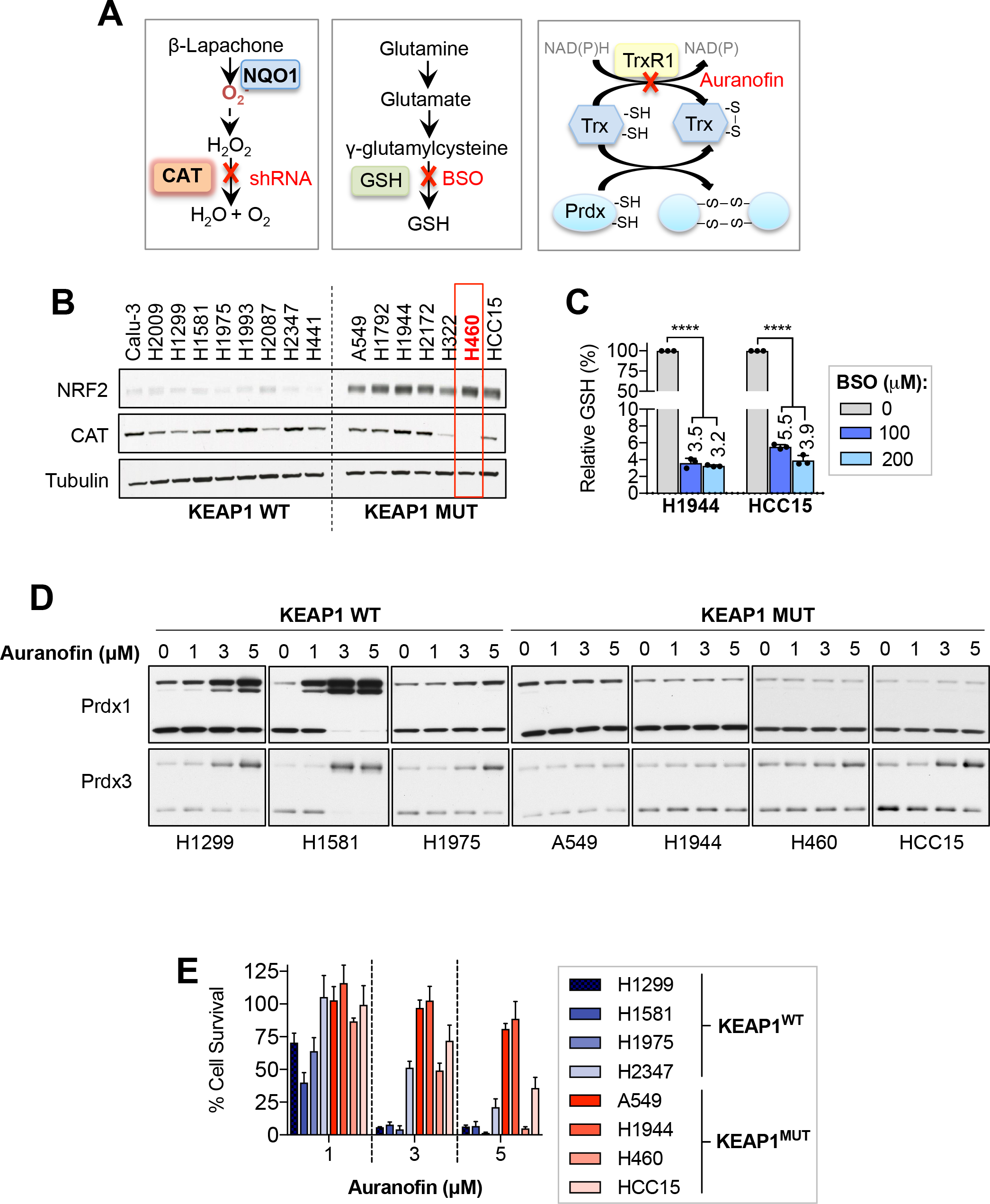
Pharmacological approaches to inhibit the TXN and GSH-dependent pathways. **(A)** Schematic representation of the role of the hydrogen peroxide scavenging systems and the compounds used in this study to specifically block these pathways. **(B)** Western blotting analysis to evaluate levels of NRF2 and catalase across a panel of KEAP1^WT^ and KEAP1^MUT^ NSCLC cells. Tubulin is included as a loading control. **(C)** Evaluation of the relative levels of reduced glutathione in cells exposed to 100 or 200 µM of BSO for 24 hours using the GSH-Glo glutathione assay (Promega). **(D)** Redox western blot analyses of evaluate the oxidation state of peroxiredoxin 1 (cytosolic) and Peroxiredoxin 3 (mitochondrial) to test the efficacy of auranofin inhibiting the TXN-dependent system. Of note, the lower band corresponds to the monomeric form of peroxiredoxins (reduced state) and the upper band results from the dimerization of peroxiredoxins (oxidized). **(E)** Survival assays of KEAP1^WT^ and KEAP1^MUT^ cells exposed to the indicated concentrations of Auranofin.

**Supplementary figure 4.**
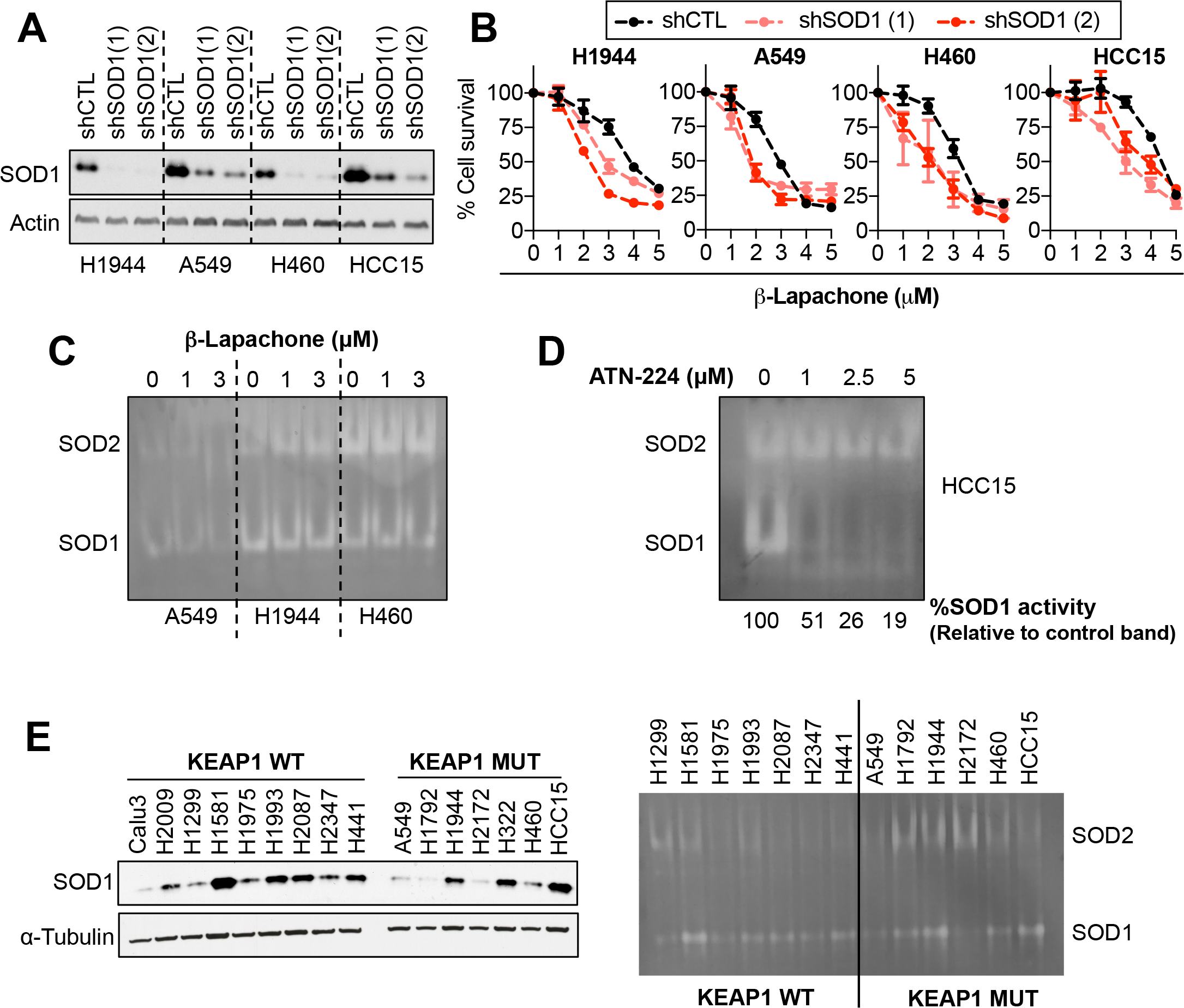
Genetic and pharmacological approaches to inactivate of SOD1. **(A)** SOD1 knockdown efficacy was validated by western blotting. Actin was included as a loading control. **(B)** Cell viability assays of KEAP1 mutant NSCLC cell lines infected with an empty vector (shCTL) or with shRNAs against SOD1 expression. Cells were treated with vehicle (0.012% DMSO) or with the indicated concentrations of β-lapachone for 2 hours. Cell viability was assessed 48 hours after treatment. **(C)** SOD in-gel activity assay of NSCLC cells exposed to 1 or 3 µM of β-lapachone or vehicle. **(D)** SOD activity gel assay of HCC15 cells treated with vehicle (0.025% DMSO) or with the indicated concentrations of ATN-224 for 24 hours. SOD1 activity was quantified by densitometry using ImageJ. **(E)** Western blotting (left) and SOD in-gel activity assay (right) of a panel of NSCLC cells to examine the SOD1 protein levels and activity across cell lines.

